# MyGene.info: gene annotation query as a service

**DOI:** 10.1101/009332

**Authors:** Chunlei Wu, Adam Mark, Andrew I. Su

**Author notes:** Correspondence may also be addressed to Chunlei Wu.

## Abstract

Biomedical knowledge is often represented as annotations of biological entities such as genes, genetic variants, diseases, and drugs. For gene annotations, they are fragmented across data repositories like NCBI Entrez, Ensembl, UniProt, and hundreds (or more) of other specialized databases. While the volume and breadth of annotations is valuable, their fragmentation across many data silos is often frustrating and inefficient. Bioinformaticians everywhere must continuously and repetitively engage in data wrangling in an effort to comprehensively integrate knowledge from all these resources, and these uncoordinated efforts represent an enormous duplication of work. We previously released MyGene.info (http://mygene.info) to enable bioinformatics developers to gain programmatic access to gene annotation data through our high-performance web services. This article focuses on the updates to MyGene.info since our last paper (2013 database issue). With the completely re-factored system, MyGene.info now expands the support from the original nine species to over 14K species, covering >17M genes with >50 gene-specific annotation types. Two simple web service endpoints provides high-performance query access to all these aggregated gene annotations. The infrastructure underlying MyGene.info is highly scalable, which offers both high-performance and high-concurrency, and makes MyGene.info particularly suitable for the use cases of real-time applications and analysis pipelines.

## INTRODUCTION

The accumulation of biomedical knowledge is growing exponentially. In an effort to organize this knowledge in a computable format, many efforts seek to structure research findings as annotations on various biological entities, such as genes, genetic variants, diseases and drugs. Among them, gene annotations are probably the most fundamental. Gene annotations are commonly stored in various biological databases. These databases include large, centralized data repositories, e.g., NCBI Entrez Gene (1), Ensembl (2), UniProt (3). In addition, important annotations also exist in more specialized resources. For example, 181 databases were listed in recent 2014 Nucleic Acids Research Database Issue alone (4). Many of them are related to genes, ranging from transcriptional factor databases and protein-interaction databases to 3D structure databases. Since biomedical research is an iterative process in which new hypotheses are based on existing knowledge, efficient retrieval and integration of these annotation data across all these resources is extremely important when formulating new hypotheses.

We previously developed an extensible and customizable gene portal system, called BioGPS (http://biogps.org), to provide a unified interface for biologists to aggregate and navigate through their relevant gene-centric annotation resources (5,6). MyGene.info was initially developed as the backend of BioGPS for hosting aggregated gene-centric annotation data. We then later released MyGene.info as public web services for general research community (5). Since our last paper was published (5), we have completely re-factored MyGene.info with a significant boost of performance and extensibility. As the result, v2 web service **API** (Application Program Interface) was released in July 2013. MyGene.info currently serves gene-centric annotations for almost 17M genes from over 14K species (compared to only nine species in the initial release). More than 50 different types of annotation data were loaded from a variety of resources, including both large data repositories (e.g. NCBI, Ensembl, Uniprot, UCSC) and also specific annotation resources like PharmGKB, PDB, Reactome, Wikipathways. MyGene.info now provides a simple and straightforward gene query API with high-performance and flexible query syntax.

In MyGene.info, all annotations relevant to the same gene are stored in a single, unified gene object, represented in **JSON** (JavaScript Object Notation) format. Different annotation types are represented as “fields” within the gene object, and each field can host data in their own data structure (See supplementary S1 for a full example of gene objects). When new annotations are available, they will be appended to the same gene object as new fields. The powerful query engine we built allows users to query any field within an object and also retrieve any subset of fields that user needs. This strategy greatly simplifies the complexity of our underlying database, as each gene annotation object is self-contained and can be updated individually when needed. This property, in turn, provides the base for MyGene.info’s high-performance and high-scalable query interface with the most up-to-date gene annotations.

## SIMPLE QUERY API

MyGene.info provides two simple-to-use REST-based web services: a gene query service and a gene annotation service. The gene query service allows users to query for gene annotations using any identifier or keyword they need, while the gene annotation service provides a convenient way to retrieve genecentric annotations when gene IDs (NCBI Entrez gene IDs or Ensembl gene IDs) are available (Figure 1).

**Figure 1.**
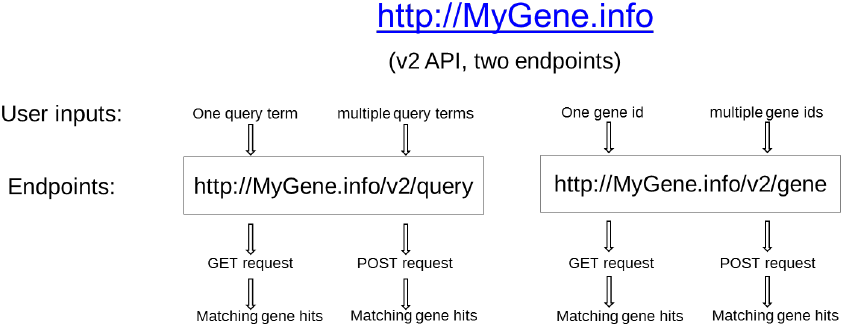
MyGene.info provides two simple-to-use REST-based web service endpoints in its v2 API: gene query service and gene annotation service.

### Gene query service

MyGene.info’s gene query service (***/v2/query***) offers a rich set of query syntax suitable for a variety of use cases (Table 1). Users can query for matching gene annotation objects by various criteria, e.g. keyword search, wildcard search, exact symbol/accession matching, genomic ranges, etc. User queries can be searched against all indexed fields or any particular field by prefixed with the field name (see “fielded queries” examples in Table 1). More complicated queries can be constructed by combining multiple query terms with Boolean operators, or conducting faceting or aggregation for special use-cases. Common query features like sorting and paging are also supported. More information on the types of queries and detailed usage instructions can be found from the documentation of MyGene.info (http://docs.mygene.info).

**Table 1.** MyGene.info gene query service supports rich query syntax. Typical query types and examples are listed in this table. More details can be found from MyGene.info documentation (http://docs.mygene.info).

| Query types | Examples |
| --- | --- |
| simple query | `/v2/query?q=cdk2` |
| customizing returned fields | `/v2/query?q=cdk2&fields=name,symbol,refseq,uniprot` |
| limited by species | `/v2/query?q=cdk2&species=human` (by common names) |
|  | /v2/query?q=cdk2&species=9606,10090,10116 (by taxonomy ids) |
| fielded queries | /v2/query?q=entrezgene:1017<br>/v2/query?q=symbol:cdk2<br>/v2/query?q=refseq:NM_001798<br>/v2/query?q=_exists_:pfam (return gene hits with “pfam” field) |
| wildcard queries | /v2/query?q=CDK?<br>/v2/query?q=symbol:CDK?<br>/v2/query?q=IL*R |
| genome interval queries | /v2/query?q=chrX:151073054-151383976&species:9606<br>/v2/query?q=chrX:151,073,054-151,383,976&species:human |
| Boolean operators and grouping | /v2/query?q=tumor AND suppressor<br>/v2/query?q=CDK2 OR BTK<br>/v2/query?q="tumor suppressor" NOT receptor<br>/v2/query?q=(interleukin OR insulin) AND receptor |

The gene query service returns a list of matching gene objects, together with their matching scores (Figure 2). By default, each gene object returns basic annotations like gene ID, symbol, name, taxonomy ID (equivalent of passing “*fields=entrezgene,symbol,name,taxid*” as a parameter). Users can specify their own “*fields*” parameter to return any combination of relevant annotations, among over 50 available annotation types, e.g., “*fields=name,symbol,refseq,uniprot*”. The complete list of available fields can be found from “*available_fields*” section of http://mygene.info/metadata (also in Supplementary file S2) and from the MyGene.info documentation (http://docs.mygene.info). To access nested fields (e.g. “uniprot” field in Supplementary file S1), “dot notation” can be passed to the “*fields*” parameter as well. For example, passing “*fields=uniprot.Swiss-Prot*” will return only Swiss-Prot UniProt IDs, while “*fields=uniprot*” will include both Swiss-Prot and TrEMBL UniProt IDs. If “*fields=all*” is passed, all available annotations for the given genes will be returned. Without specifying a “*species*” parameter, the matching gene hits from human, mouse and rat are returned as default. Users can customize the returned hits by passing one or multiple species (either common names or their taxonomy ids, out of over 14K supported species) to “*species*” parameter.

**Figure 2.**
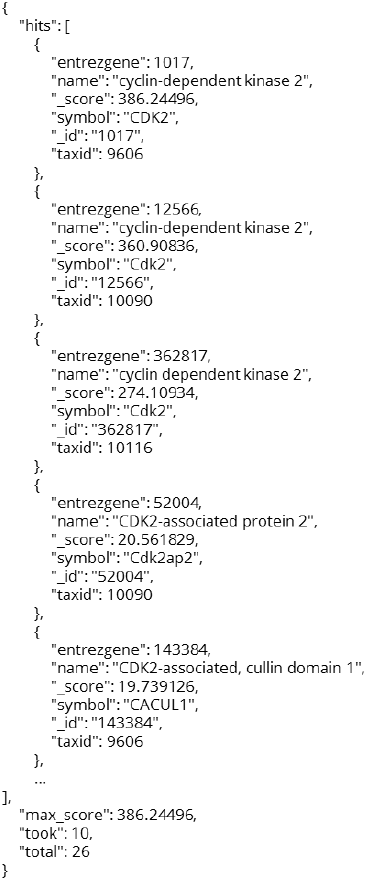
An example of returned gene hits, in JSON format, from MyGene.info gene query service. The gene hits are sorted by their matching scores by default. Users can customize the returned annotation fields for each gene object by specifying “fields” parameter.

In common use cases, users can access the gene query service via standard HTTP GET request with their desired query term. When users have a list of query terms, e.g., a list of gene symbols (hundreds or even thousands) from an analysis pipeline, it’s often more efficient to submit queries in batches. The gene query service supports batch queries with HTTP POST request on the same service end-point. Users can pass a list of query terms, and also specify one or multiple field names as the “*scopes*” parameter, which will limit the query scopes of the input query terms, e.g., symbols, UniProt IDs, RefSeq accessions. As “scopes” parameter supports multiple fields, users’ input query terms can have mixed types. For example, it’s not uncommon for bioinformaticians to handle a list of sequence IDs mixed of UniGene, RefSeq and GeneBank IDs. In that case, passing “scopes=unigene,refseq,accession” will map all these IDs into matching genes in one query. By combining with the “fields” parameter, batch mode of gene query service is particularly suitable for routine ID translation tasks, even with messy input.

### Gene annotation service

MyGene.info’s gene annotation service (***/v2/gene***) takes a canonical gene ID (either a NCBI Entrez gene ID or an Ensembl gene ID) and returns relevant annotations for that particular gene. It is essentially a convenient wrapper of gene query service where the input is always gene IDs. Just like gene query service, it also takes a single gene ID via GET request, and a list of gene IDs via POST request for batch mode (Figure 1). The same “fields” parameter can be used to customize returned types of annotations.

When an input gene ID is not valid, an HTTP 404 (NOT FOUND) response will be returned. It’s worth noting that retired Entrez gene IDs are also supported by gene annotation service. For example, gene “*245794*” is retired and replaced by gene “*10015*”, so the request to “*/v2/gene/245794*” will return the annotation object of gene “*10015*” instead, with the “*retired*” field as “*245794*”. However, if a retired gene ID is completely discontinued without any replacement, this gene ID is no longer valid and 404 will be returned.

### Metadata service

Besides the gene query and annotation services which provide the access to actual gene annotations, we also provide a metadata service (**/metadata**) for users to access the metadata about the annotations hosted at MyGene.info, such as the gene statistics, the list of available fields, supported species common names, the versions/timestamps of all data sources we loaded the data from (See supplementary file S2 for an example of metadata output).

## ALWAYS UP-TO-DATE DATA BACKEND

Gene annotation data can be very broad and heterogeneous due to the diverse nature of the data and the data sources. In MyGene.info, we represent a gene annotation object in JSON format. Originally developed for use in web applications, JSON is a lightweight data-interchange format that has quickly gained popularity in general applications as well. JSON is concise and human-readable, while still remaining machine-readable. It also has rich data structure support, and includes almost everything used in modern programming languages, such as records, lists and trees. This makes JSON particularly suited for representing heterogeneous gene annotation objects. All annotations related to the same gene can be incorporated into the respective individual objects, while maintaining their own preferred structures. Additionally, a JSON object can be easily extended by adding new annotation fields without breaking accessibility to existing fields.

To build an aggregated annotation object, the first step is to convert each individual annotation data into individual JSON objects. Different annotation type can have its own data structure, and the only requirement is that each individual object contains an “*_id*” field as the primary key. That way, we can then merge all gene-specific annotations together based on the primary key. Gene IDs are the natural choice for the primary key. Gene IDs from NCBI’s Entrez Gene and Ensembl are the two most commonly used gene ID systems in biological research. We arbitrarily picked NCBI’s Entrez gene IDs as the preferred primary keys. That is, when an Ensembl gene ID can be mapped to a corresponding Entrez gene ID, we use Entrez gene ID as the primary key. It’s worth noting that Ensembl gene ID is still accessible and queryable from the JSON object as a field, just not chosen as the primary key. In the case that an Ensembl gene ID cannot be mapped to an Entrez gene ID (e.g. a unique gene identified by Ensembl), we then use the Ensembl gene ID as the primary key.

By following this simple merging rule, we wrote an individual data parser for each source annotation data. Each parser can have its own update schedule. The output of each parser, a list of JSON objects, is stored in a MongoDB (http://mongodb.org) instance with timestamps recorded. Then the merging algorithm takes the latest version of each individual annotation objects and combines all objects with the same primary key (“*_id*” field) together into a single annotation object (Figure 3). The advantage of this setup ensures that the processing of each annotation source is independent with others, at their own update schedules, and any single failure will not cause the entire merging process to fail. In the case that a single parser fails, the last success version of that failed source will be used, while the parser is being fixed.

**Figure 3.**
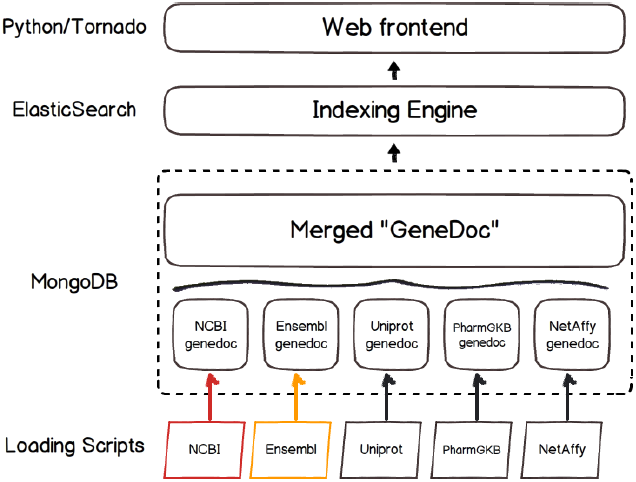
Schematic design of the MyGene.info architecture. Colors depict different update frequencies.

This architecture drastically increases the efficiency of data management and allows MyGene. Info to remain in nearly real-time sync with the source databases. Currently, the data aggregation backend of MyGene.info loads all available genes from both NCBI Entrez and Ensembl (>17M genes from over 14K species) as the base gene objects (including basic annotations like gene IDs, names, symbols, etc.), and additional gene-specific annotations will be appended from varieties of resources. There are more than 50 different types of gene annotations loaded so far, and additional annotation types are continuously added over time. Our data backend supports automatic and asynchronous updates from multiple data sources, each of which can have its own update frequency. As soon as a new release is available from a data source, new data will be updated immediately at MyGene.info. For those resources with daily roll-out, e.g. NCBI Entrez dumps, they are updated at least weekly.

After the merging process, each gene object contains all aggregated annotations for a given gene. We then built a powerful Elasticsearch (http://elasticsearch.org) based query engine to index all fields within these annotation objects, so that users can query for gene hits matching their search criteria or retrieve relevant annotations for their input genes (Figure 3). Elasticsearch is a highly scalable open-source full-text search and analytics engine. It provides a rich query syntax and inherent scalability to handle large-scale data loads in real time. On top of Elasticsearch, we built user-friendly REST web service APIs using Tornado web framework (http://tornadoweb.org). Tornado is a Python-based web framework built upon asynchronous networking technology, which can provide tens of thousands of concurrent connections with a moderate server.

## HIGH PERFORMANCE WEB SERVICES

Since the release of the v2 API (July 2013), MyGene.info has accumulated over 70 million requests, and currently serves more than 2 million requests per month. Approximately 40% of traffic comes from our BioGPS application, while 60% of traffic comes externally from ∼3000 unique IP addresses. Figure 4 shows the recent 30-day request counts with the peak value around 30,000 requests per hour; and Figure 5 shows the majority of the actual user requests take less than 30ms.

**Figure 4.**
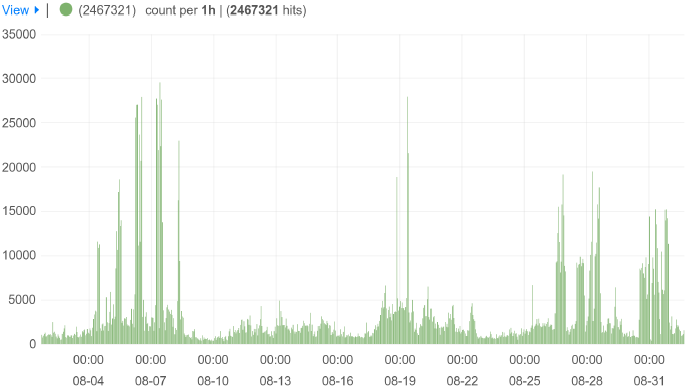
Hourly request counts for MyGene.info services during recent 30-day period (08/01/2014-09/01/2014). The peak values are around 30,000 requests per hour.

**Figure 5.**
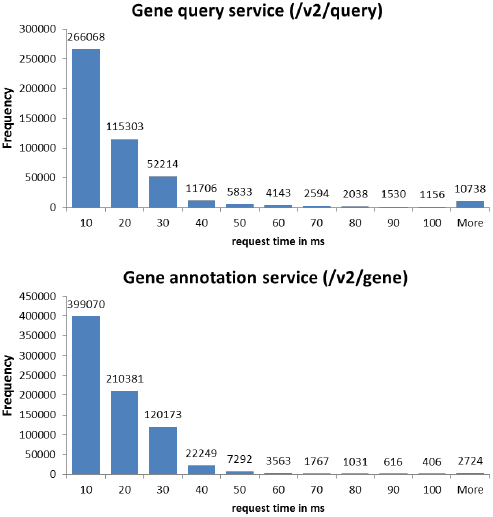
Histograms of the request time in mille-second for actual user requests to the gene query service (up) and the gene annotation service (down). The number on each bar indicates the actual count. The data were extracted from the logs on one server node during recent 30-day period (08/01/2014-09/01/2014).

The performance, scalability and stability are three key features of a successful web-service provider. The MyGene.info system is currently hosted on Amazon EC2 platform with four moderate servers at an estimated cost of $200 per month. At current usage, MyGene.info receives less than 500 requests per minute, however, our internal stress tests indicate that the current setup can achieve the performance capacity of >5,000 concurrent users for ∼10,000 requests per minute. Both Elasticsearch and Tornado can be linearly scaled by adding identical nodes into the system to meet the increasing scalability demands as usage grows. Upon system upgrade or failure, individual nodes can be shut down or added at any time without causing downtime to the services. As the result of this general design, MyGene.info achieved 100% uptime since its v2 release in July 2013 (http://bit.ly/mygene-info-uptime). We are very confident that we can provide robust, high-performance and high capacity web services to the biological research community, allowing the use of remote service calls for real-time annotation data queries.

## CLIENTS AND WIDGETS

### Python Client

With the previous v1 API, we released a Python client, called ***mygene.py*** (v1.0.0), for even tighter and easier integration in Python applications. We have now updated it (currently at v2.2.0) to utilize the high-performance v2 API, and also expanded the support for newer Python3. Currently, *mygene.py* module receives more than 1500 downloads per month (http://pypi.python.org/pypi/mygene), contributing about 20% of total MyGene.info requests (stats based on 08/1/2014-09/01/2014).

### R Client

Along with our Python client, we recently released our R Client, called ***mygene.R***, which is currently pending approval for inclusion in Bioconductor (http://bioconductor.org). Users can already install *mygene.R* from the source (https://bitbucket.org/sulab/mygene.r). Besides providing the wrappers around the MyGene.info web API, *mygene.R* also offers tight integration with other Bioconductor packages by returning data in DataFrame, GRanges, GRangesList and TranscriptDb classes (7). Compared to existing downloadable gene annotation packages commonly used in current Bioconductor, *mygene.R* provides much broader species coverage (over 14K species versus only 30 species supported by the latest Bioconductor v2.14). It is very light-weighted, provides always-up-to-date gene annotations without the need for downloading or updating local annotation packages. Additionally, *mygene.R* offers more flexible query capability, supporting complex queries across species or across multiple resources.

### Auto completion widget

Building web applications to distribute analysis tools or publish their results has been increasingly popular among bioinformatics developers. As REST-based web APIs, MyGene.info services are very easy to integrate in web applications, so that developers don’t have to maintain their own local annotation databases and worry about the routine updates over time. As a common feature, many biological web applications implement a gene search box for users to query genes by their symbols, names or other identifiers, and then display gene-specific data relevant to the applications. Built on MyGene.info services, we also provide a ready-to-use JavaScript widget for developers to plug into their applications. This widget renders a gene search box with auto-completion enabled (https://bitbucket.org/sulab/mygene.autocomplete), without any backend database or code needed. Users can then write a callback function to define the actions when a gene is selected from the dropdown list (Supplementary Figure S3). Furthermore, users have many options to customize the desired behavior, such as the search scopes (e.g. limit the query to human-only genes or ncRNA genes), display labels in the dropdown menu and the returned values when a gene is selected.

## SOURCE CODE

MyGene.info is an open source project. The source code of this project contain two components: web service front end (repository: https://bitbucket.org/sulab/mygene.info; license: Apache2) and data aggregation backend (repository: https://bitbucket.org/sulab/mygene.hub; license: GNU GPLv3)

## DISCUSSION

The current fragmentation of biological annotations results in huge inefficiencies in biomedical research. First, every bioinformatician has written data parsers in an effort to integrate data from multiple resources, and these uncoordinated efforts represent an enormous duplication of work. Second, these individual efforts vary widely in terms of both completeness and accuracy, affecting the quality of downstream experiments on which these analyses are based. And third, these resources quickly go out of date because the landscape of annotation resources is continuously changing.

The solution provided by MyGene.info essentially represents a centralized database with web services as the interface for data access. Web services provide an elegant way for users to access structured data remotely, by keeping requirements for client-side dependencies very low. It has been increasingly popular for biological database providers to provide data access via web services, for example NCBI eUtils (8–11), BioMart (12) and UniProt web services (3). There are also efforts to make aggregated annotation databases across multiple data sources with web service access (13,14). Compared with those existing resources, MyGene.info distinguishes itself in these respects: a) broad annotation and species coverage; b) superior query performance with the support of high concurrency; c) Simple API (just two service endpoints for all query cases), but with great query flexibility; d) tight weekly updates to deliver always up-to-date annotations.

Finally, although MyGene.info is a gene-specific annotation resource, its underlying architecture is suitable for aggregating and serving annotations for other biological entities as well, such as genetic variants, disease and drugs.

## Acknowledgement

The authors acknowledge contributions from Ryan Thompson for building mygene.R client and valuable feedback from the MyGene.info user community.

